# Shared genetic etiology of age of menarche and socioeconomic variables: No evidence for genetic overlap with psychiatric traits

**DOI:** 10.1101/2020.05.01.072348

**Authors:** Martin Steppan

## Abstract

Earlier research has shown observational associations of early pubertal timing and poor mental health. Mendelian randomization (MR) studies demonstrated a transient effect of pubertal timing on mental health during adolescence, but not later in life. MR studies also showed that there is a likely causal association of pubertal timing with life history traits. However, the strongest causal effects and genetic correlations with age of menarche have been found for Body Mass Index (BMI). As high BMI is associated with lower socioeconomic status and with poor mental health, the shared genetic etiology of socioeconomic status, BMI and poor mental health is not yet fully understood. BMI correlates negatively with socioeconomic status and several mental health outcomes. Despite their substantial genetic overlap, the underlying genetic etiology of these phenotypes remains unclear. In this study we applied Linkage Disequilibrium score regression to test genetic correlations of age of menarche with 33 socioeconomic, life history, social interaction, personality and psychiatric traits, and BMI. We further applied spectral decomposition and hierarchical clustering to the genetic correlation matrix. After controlling for multiple testing, we could only identify significant genetic correlations with BMI and three socioeconomic traits (household income, deprivation and parental longevity). The results suggest that genome-wide association studies on age of menarche also contain socioeconomic information. Future MR studies aiming to test the unconfounded effect of pubertal timing should make sure that genetic instruments have no pleiotropic effect on socioeconomic variables, or (if possible) also control for socioeconomic status on the observational level.

## 1 Introduction

Previous research focussing on age of menarche revealed that early pubertal timing is a potential risk factor for the development of somatic and mental health problems [1, 2]. To date, several studies have investigated genetic correlations of and causal effects on age of menarche using Mendelian Randomization (MR). The most striking finding was the association between Body Mass Index (BMI) and age of menarche. In their paper introducing the technique of LD Score Regression [3], Bulik-Sullivan et al. had already included age of menarche and its genetic correlations with 23 other traits, of which only body mass index, childhood obesity (both negative) and height (positive) showed significant genetic correlations with age of menarche. MR-studies have demonstrated the causal association of BMI and pubertal timing in both directions, suggesting a causal effect of BMI on pubertal timing and vice versa [4, 5]. In a recent paper, a genetic correlation was also found for male pubertal timing and BMI [6].

However, there is no evidence on genetic correlations of age of menarche with psychiatric disorders. The earlier mentioned study by Bulik-Sullivan et al. included genetic correlations with psychiatric disorders (Autism, depression, anorexia, bipolar and schizophrenia), of which none showed a significant genetic correlation with pubertal timing [3]. Moreover, a recent study tested genetic correlations of antisocial behavior with age of menarche, but did not find a significant genetic correlation [7]. In a study on male puberty, no significant genetic correlations were found with psychiatric problems [6]. Nonetheless, studies based on Mendelian Randomization [MR] did show a transient effect of age of menarche on mental health [depressive symptoms] [8], but not later in life [9]. Hence, these studies indicate a likely transient causal link, but no shared genetic etiology of pubertal timing and mental health.

Having said this, all three domains - age of menarche as well as BMI and mental health - seem to be confounded with socioeconomic status. For example, a recent MR study showed a significant causal effect of age of menarche on traits like educational attainment, age at first and last birth, childlessness and alcohol consumption [10], indicating socioeconomic effects. Bulik-Sullivan et al. had already reported a genetic correlation of age of menarche with educational attainment and BMI [3], suggesting shared genetic architecture underlying pubertal timing and socioeconomic variables and BMI. A recent MR-study has also found likely causal effects of BMI on poor mental health and well-being [11]. Hence, it is not surprising that a recent study investigating 751 genetic correlations of male pubertal timing, found positive genetic correlations with maternal longevity, intelligence, an overall health rating, education and a strong negative genetic correlation with BMI [6].

In sum, age of menarche, socioeconomic traits, BMI and mental health showed genetic overlap and causal effects on each other in a variety of studies. These findings suggest that socioeconomic status is a potential confounder in this framework, potentially affecting associations of biological relevance, due to an ecological clustering of certain health and psychological problems within social milieus. Therefore, the aim of this study is to elucidate the interplay of genetic correlations among these four domains. We aim at achieving this by adding additional psychiatric problems that have not been investigated yet, personality traits and additional socioeconomic traits, to better understand the genetic overlap of these variables. Moreover, the analyses fulfills the purpose to allocate pubertal timing within the larger space of genetic correlations using spectral decomposition. Lastly, we will explore whether genetic instruments reported by Perry et al. [12] can be used in a follow-up MR-study, or whether additional control of socioeconomic confounding needs to be undertaken.

## 2 Methods

We used cross-trait linkage disequilibrium score regression to estimate genetic correlations between pubertal timing (age of menarche) and 33 socioeconomic, reproductive, psychological and psychiatric traits, as well as body mass index.

### 2.1 Ethical approval

This study is based on publicly available summary data from thirtythree genome-wide association studies (GWAS). No individual data was analysed in this study. Hence, no ethical approval was required.

### 2.2 Sample

Table 1 shows the genome-wide association studies, which we included and for which summary data can be accessed online. Sample sizes varied between 4596 and 2,007,916 individuals. Seventeen out of 33 traits were continuous, i.e. there was no distinction between cases and controls. All of the used samples were of European ancestry. For pubertal timing we used those summary statistics which excluded individuals from a collaborative oncological gene-environment study [12]. We could not find publicly available summary statistics for male pubertal timing. However, Hollis et al. [6] tested genetic correlations of male pubertal timing with several of the traits used in this study. The authors also report a genetic correlation of male and female pubertal timing of *r*_*g*_ = .683, p<2.6E-213, indicating that genetic correlations of male and female pubertal timing should show substantial agreement.

**Table 1:**
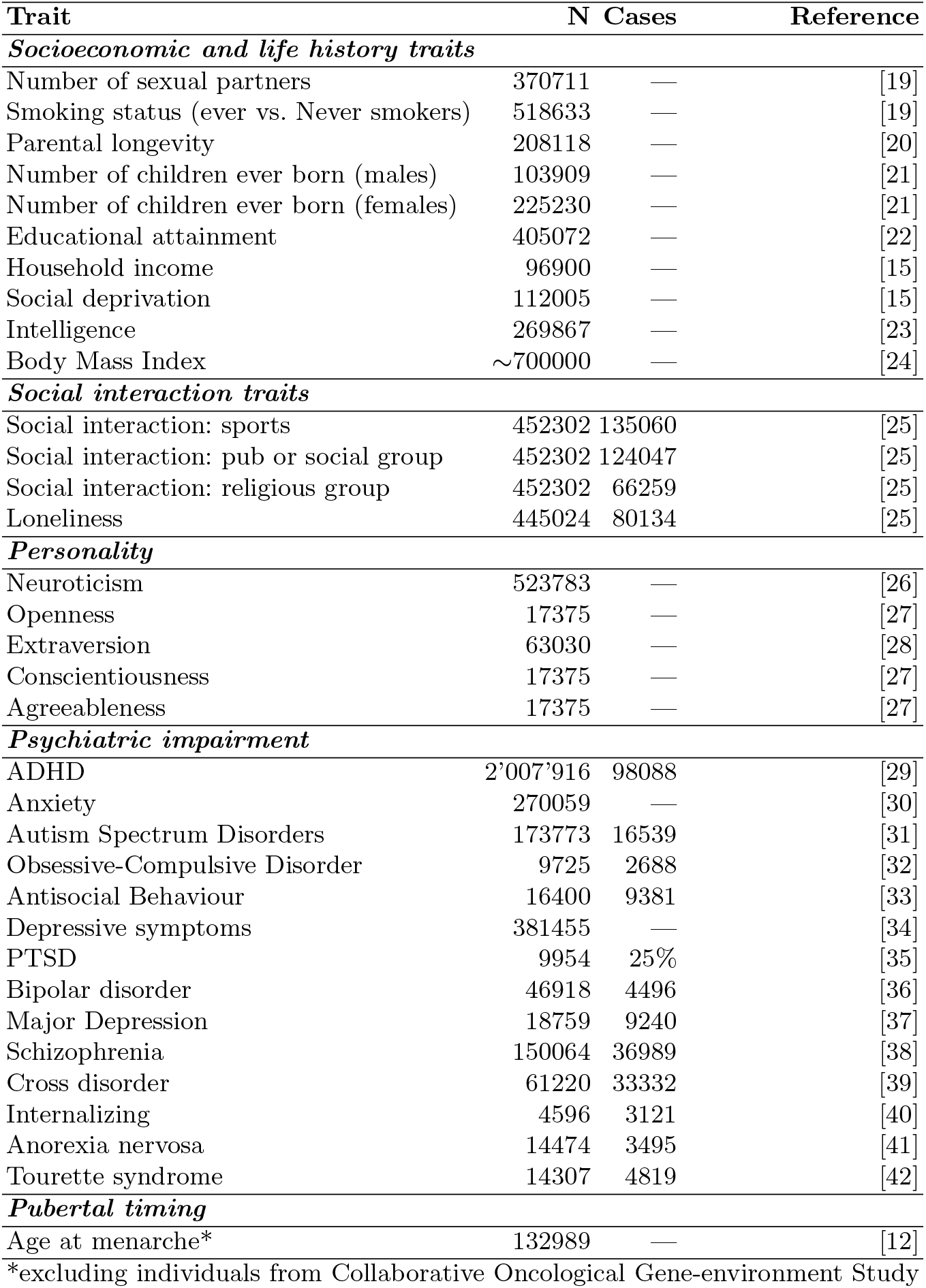
Included genome-wide association studies

### 2.3 Statistical analysis

We applied a python script authored by Bulik-Sullivan and Finucane. The matrix of genetic correlations was visualized using the R package “corrplot”. [13]. A reordering of the traits was based on hierarchical clustering [“hclust”]. The first three clusters were visualized in the correlation plot. Spectral decomposition of the correlation matrix was performed using the R base function “eigen”.

## 3 Results

Genetic correlations revealed that pubertal timing is genetically related to body mass index and socioeconomic traits, but not to personality and psychiatric impairment. We found genetic overlap of pubertal timing with body mass index, parental longevity, deprivation, household income and anxiety. Figure 1 illustrates all genetic correlations with age of menarche, their direction, p value and to which category they belong. The strongest overlap was found for body mass index (*r*_*g*_=−.33). Socioeconomic traits showed equivocal genetic correlations in the sense that early puberty was genetically correlated with indicators of lower socioeconomic status (i.e. higher deprivation, lower household income). Parental longevity also follows this pattern, i.e. early puberty is genetically correlated with lower parental longevity. There were no significant genetic correlations of pubertal timing with personality or social interaction traits. Except for anxiety, there were also no significant genetic correlations with psychiatric variables. The genetic correlation with anxiety, however, was not significant when controlling for all tests (p<.05 / 528), and was only significant when controlling for all tests with age of menarche (p<.05 / 33). The result for anxiety may also be related to a significant genetic correlation between BMI and anxiety (see figure 2). SNP heritability of age of menarche was 0.1649 (SE: 0.0081), Lambda GC: 1.2531; Mean Chi Squared: 1.406; Intercept: 0.9963 (SE: 0.0109).

**Fig. 1:**
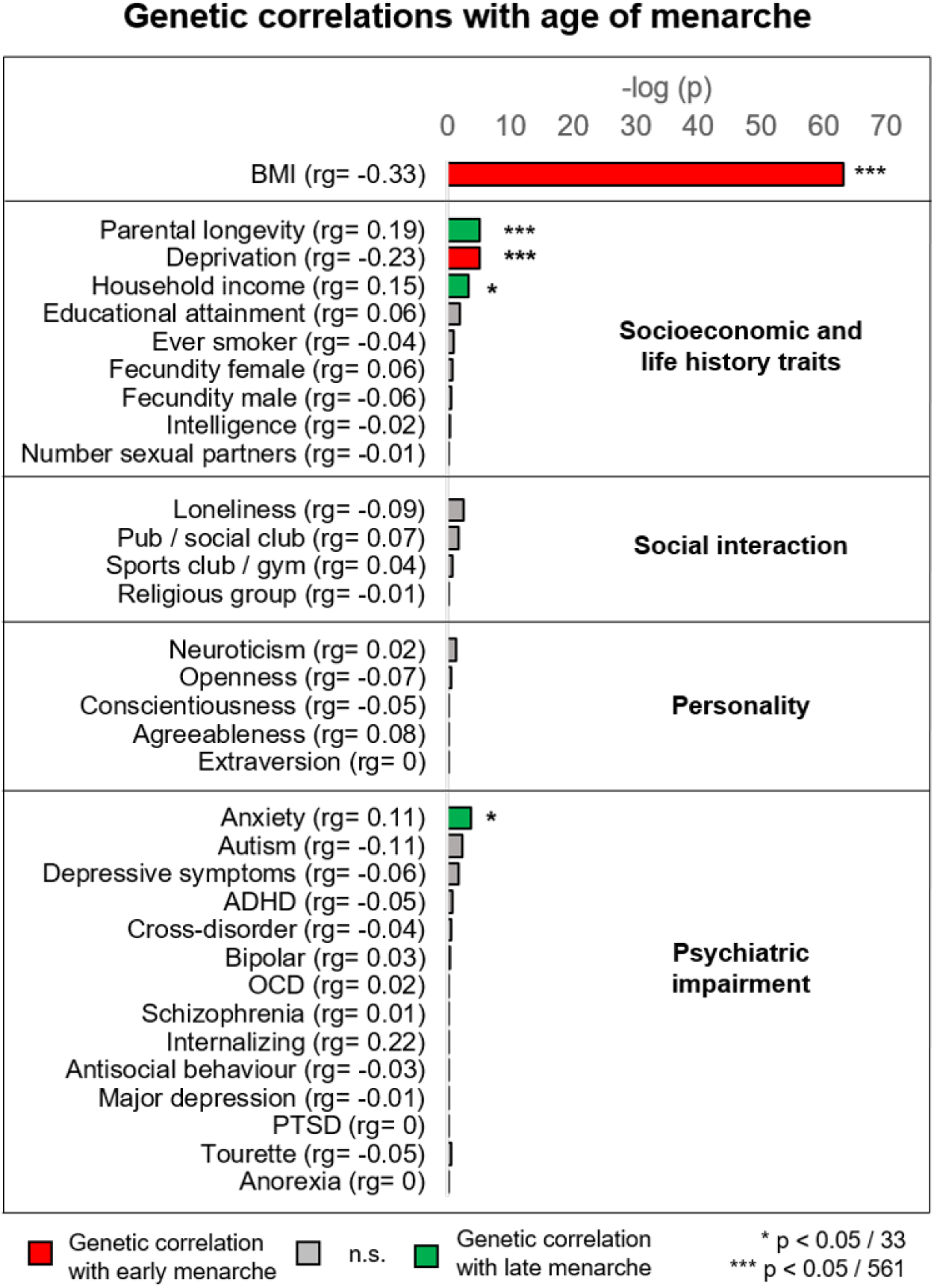
Genetic correlations of age of menarche with different socioeconomic and life history traits, social interaction variables, personality factors and psychiatric variables.

**Fig. 2:**
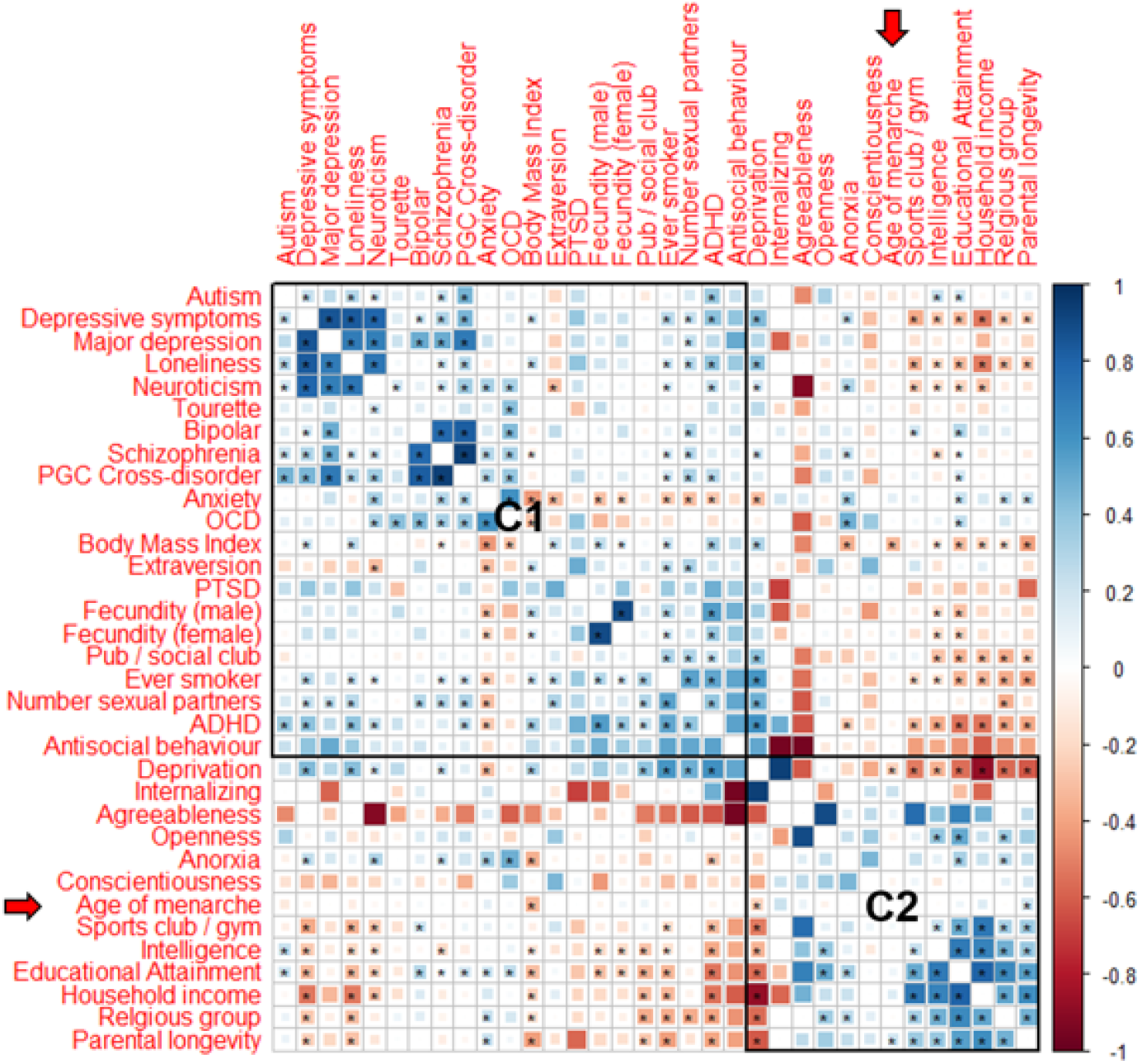
Genetic correlation matrix of all variables and their cluster membership. Significant genetic correlations (controlled for multiple testing p ¡ .05 / 561). Variables reordered based on “hclust”. C1 = cluster 1; C2 = cluster 2.

Hierarchical clustering shows that pubertal timing is closer to socioeconomic than psychiatric traits. Figure 3 shows the genetic correlation of all 33 traits and pubertal timing, and reorders the traits based on hierarchical clustering. Cluster 1 (C1) represents a cluster of psychiatric impairment. All psychiatric variables [except for anorexia and internalizing behavior] are located within this cluster. Age of menarche was classified within cluster 2 (C2), representing a cluster of socioeconomic variables and cognitive variables [educational attainment, intelligence, household income, parental longevity, but also social interaction in sports clubs or religious groups].

**Fig. 3:**
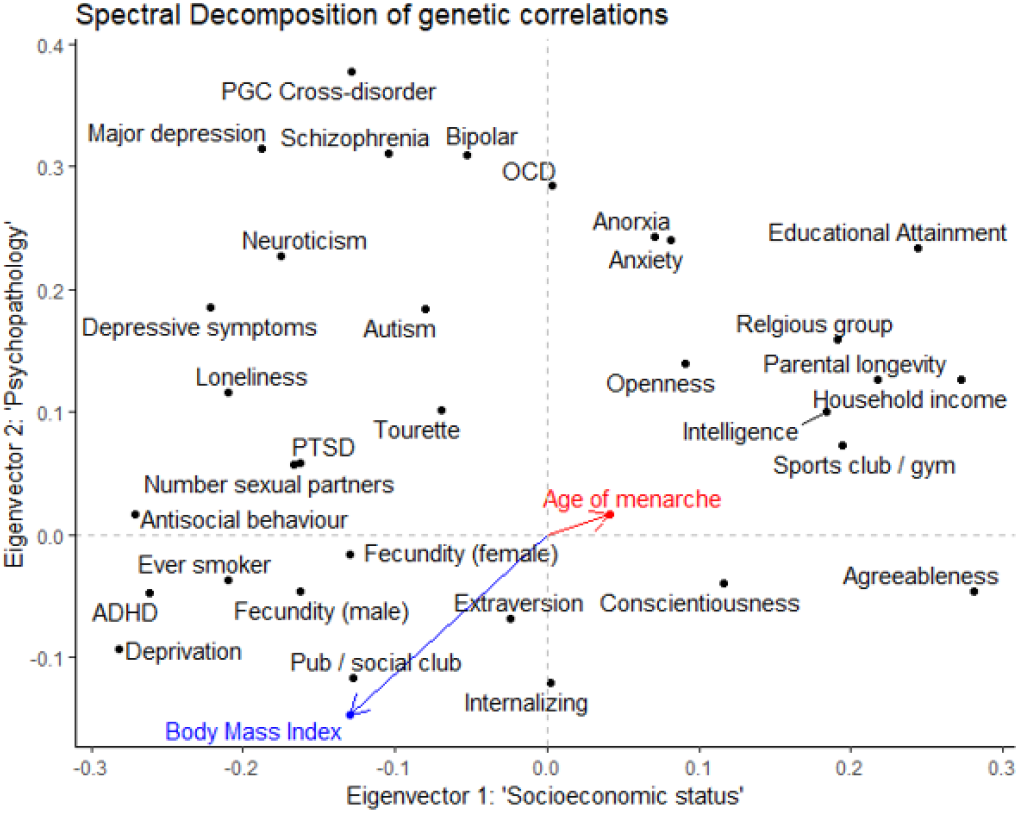
Spectral analysis (first two eigenvectors) applied to all genetic correlations (age of menarche, psychiatric diagnoses, personality factors, social interaction, socioeconomic and life history traits)

Spectral analysis (Figure 3) revealed that pubertal timing [age of menarche] does not show a substantial loading on a psychopathological eigenvector [y-axis; Eigenvector 2], but only a small negative loading on a socioeconomic eigenvector (x-axis; Eigenvector 1). Figure 3 corroborates the results that pubertal timing represents a trait with slight genetic overlap with socioeconomic features, but practically none with psychiatric impairment. The analysis also shows that some psychiatric problems are more associated with higher socioeconomic status [e.g. anorexia and anxiety]. This may explain partly why late pubertal timing show some evidence to be genetically correlated with higher anxiety (Figure 2).

## 4 Discussion

We demonstrate that pubertal timing shows a shared genetic architecture with body mass index and socioeconomic traits. There is practically no genetic overlap with psychiatric impairment, personality and social interaction variables. Only anxiety seems to be genetically related to late pubertal timing (when applying less conservative control for multiple testing). However, the association was less strong, and may be related to the strong genetic correlation of anxiety and body mass index. The results suggest that summary statistics of genome-wide association studies on pubertal timing (e.g. those analyzed here) contain socioeconomic information, potentially through the social determination of body mass index. Considering that Mendelian Randomization can be confounded by genetic overlap [14), follow-up studies applying Mendelian Randomization techniques need to make sure that methods are controlled for body mass index and socioeconomic influences.

The analyses corroborate the well-established genetic correlation of pubertal timing and body mass index [3]. The results are also identical with earlier studies which tested pairs of the exact same genetic correlations, corroborating the technical accuracy of the performed analyses [7]. The results leave no doubt regarding the genetic overlap of pubertal timing and body mass index. However, we also identified genetic overlap of pubertal timing s with socioeconomic traits. It is not clear to which extent these genetic correlations indeed represent shared genetic etiology or mediated pleiotropy [15]. According to Hill et al. [15], since socioeconomic status has “no clear biological analog”, mediated pleiotropy appears more likely than biological pleiotropy. On the one hand, genetic correlations are considered powerful in detecting confounding [16], on the other hand it is indeed questionable what a biological pathway could be, how a set of genetic markers can affect socioeconomic status as well as reproductive aging. Since most of the socioeconomic traits tested here, also showed significant genetic correlations with body mass index (see figure 2), a likely explanation is that the genetic correlations of socioeconomic traits with age of menarche are driven by their genetic overlap with body mass index. Future research might elucidate the nature of these genetic correlations using more complex multivariate approaches [14]. The current study benefits from earlier genome-wide association studies of which some had very large sample sizes.

The methodology applied (LD score regression), although widely adopted, has strengths and limitations: (a) the GWAS we included were only of European ancestry, which increases the comparability on the one hand, but also reduces the generalizability of the results. Genetic correlations may be substantially different in other populations. (b) 41 genetic correlations exceeded the normal range of genetic correlations [−1,+1]. This problem is well known, and can be a consequence of large standard errors. Nonetheless, we set these genetic correlations missing, so that they did not bias the results. (c) The sample sizes of different GWAS we included, varied considerably. Assuming that GWAS get more precise as the sample size increases, also the genetic correlations here are not estimated at the same level of precision. Hence, we assume more imprecision in particular for psychiatric diagnoses, where sample sizes were comparably low (e.g. PTSD, internalizing problems). (d) We applied spectral decomposition to the genetic correlation matrix. There is evidence that the orientation of eigenvectors can be unstable [17]. Nonetheless, we found highly meaningful eigenvectors, i.e. socioeconomic traits and psychiatric diagnoses loaded on separate eigenvectors, corroborating the validity of these results. (e) It has been shown that observational and genetic correlations highly correspond, also known as “Cheverud’s conjecture” [18]. Given this strong association, it is not clear yet, under which circumstances genetic correlations deviate from observational correlations, and what the added value of genetic correlations really is.

Future research might benefit from these results. In terms of genetic correlations, pubertal timing in girls appears much more a socioeconomic than a psychologically relevant trait. Considering the fact that summary statistics from genome-wide studies on age of menarche contain socioeconomic information, future studies applying Mendelian Randomization need to disentangle the biological from the socioeconomic signal in their instruments, in order to provide estimates which are not confounded by milieu-specific effects.

**Table 2:**
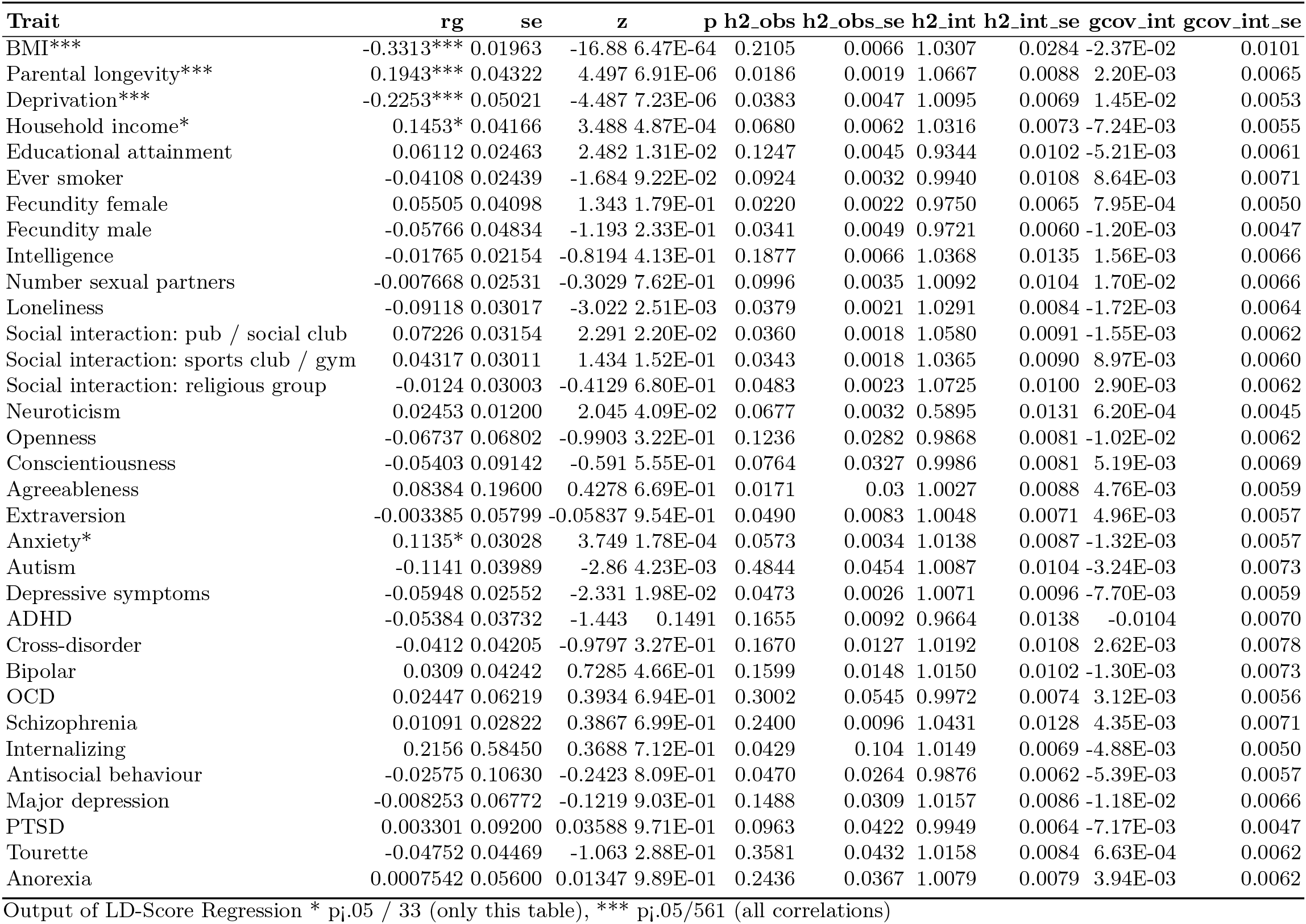
Detailed results of Linkage Disequilibrium (LD) Score Regression of age of menarche against 33 other traits

## 5 Declaration of interests

The author(s) declare no competing interest.

## 6 Acknowledgement and Funding

This study was part of a funded research project of the first author at the University of St Andrews, School of Medicine. Many thanks to Jorim Tielbeek from the University of Amsterdam for discussing this manuscript.

## Notes

### Competing Interest Statement

The authors have declared no competing interest.

